# Parameterization and Application of the General Amber Force Field to Model Fluro Substituted Furanose Moieties and Nucleosides

**DOI:** 10.1101/2022.01.26.477958

**Authors:** Diego E. Escalante, Courtney C. Aldrich, David M. Ferguson

**Affiliations:** Department of Medicinal Chemistry, University of Minnesota, Minneapolis, MN 55455; Center for Drug Design, University of Minnesota, Minneapolis, MN 55455

## Abstract

Molecular mechanics force field calculations have historically shown significant limitations in modeling the energetic and conformational interconversions of highly substituted furanose rings. This is primarily due to the gauche effect that is not easily captured using pairwise energy potentials. In this study, we present a refinement to the set of torsional parameters in the *General Amber Force Field* (*gaff*) used to calculate the potential energy of *mono, di-*, and *gem*-fluorinated nucleosides. The parameters were optimized to reproduce the pseudorotation phase angle and relative energies of a diverse set of mono- and difluoro substituted furanose ring systems using quantum mechanics umbrella sampling techniques available in the IpolQ engine in the Amber suite of programs. The parameters were developed to be internally consistent with the *gaff* force field and the TIP3P water model. The new set of angle and dihedral parameters and partial charges were validated by comparing the calculated phase angle probability to those obtained from experimental nuclear magnetic resonance experiments.

## Introduction

The selective introduction of fluorine or fluorine-containing functional groups to small organic molecules has proven extremely useful in modifying the physicochemical and pharmacokinetic properties of small drug molecules.^2-4^ In the field of nucleoside and nucleotide chemistry, fluorine substitutions to the 2’ and 3’ positions of the sugar moiety have been shown dramatically effect both metabolic stability and biological activity.^5-7^ This is not only due to the inherent strength of the C-F bond that is highly resistant to metabolic cleavage but also to the polarizing effect of the fluoro group that influences the sugar pucker angle through the gauche effect.^4, 7^ In fact, early work on ribo- and arabino-based nucleosides indicated that the position of the fluorine (either up-beta or down-alpha in the 2’ or 3’ position) was a primary determinant of molecular conformation of the sugar moiety which further influenced the orientation of the base.^8, 9^ Based on a combination of NMR experiments^1, 10, 11^ and ab initio molecular orbital calculations,^10, 11^ the preference for a primarily North or South pucker profile was shown to trend with the “ up-down” orientation of the fluoro substitution as shown in figure 1. Of course, the extent of this behavior is driven by the strength of the gauche effect that is known to be correlated to the electronegativity of the substituent. In the case of the 2’-fluro-2’,3’-dideoxyribose, this effect induces a more Northern pseudorotation angle that is further reinforced by the anomeric effect (for beta-nucleosides).^12^ In contrast, the analogous 2’-fluoro-2’,3’-arabinose derivative adopts a more Southern conformation.^1^ It is important to point out that anomeric effect actually opposes the gauche effect in this case, resulting in a lower barrier to pseudorotation and a greater distribution of pucker angles and conformational states.

**Figure 1.**
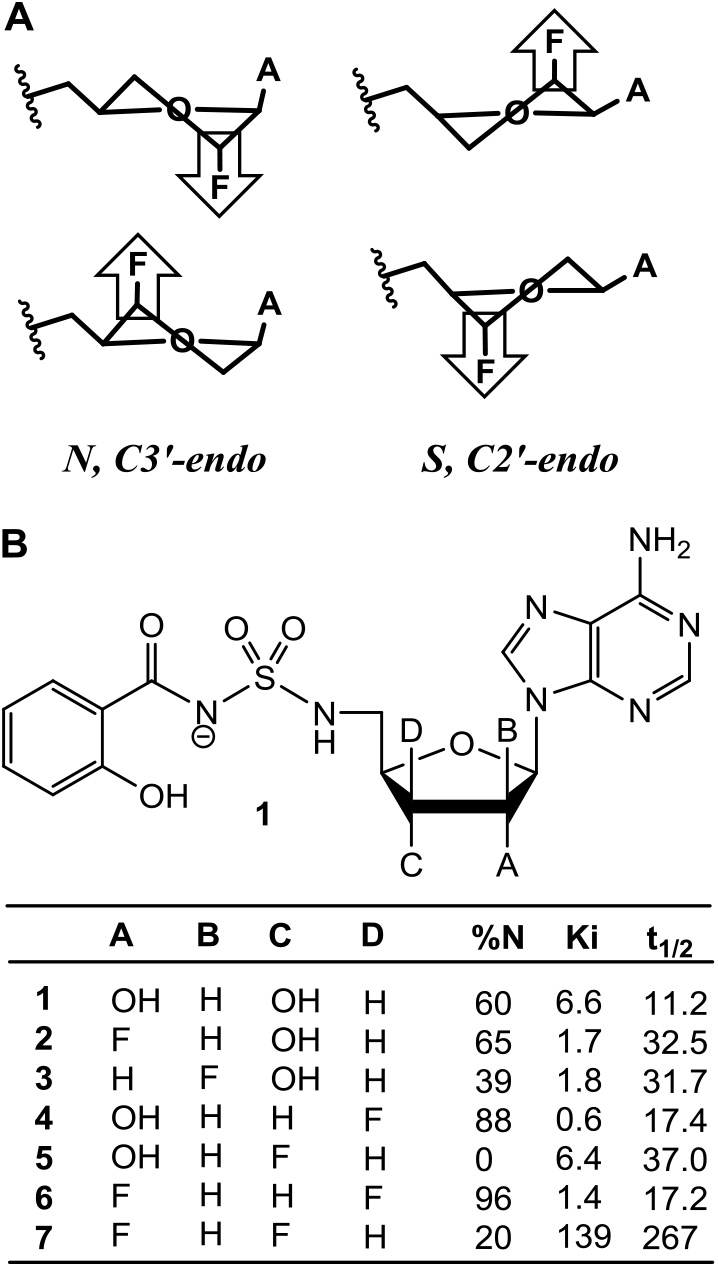
A) Influence of fluorine on ring conformation. The ring pucker is pulled to the side of the most electronegative atom due to the gauche effect. B) Ring conformation, enzyme binding, and half-life of fluorinated Sal-AMS derivatives reported in reference X.

Prior work by our lab has shown fluorination can also have a significant impact on the metabolic stability and biological activity of complex nucleosides.^1^ <u>Sal</u>icyl <u>a</u>denosine <u>m</u>ono<u>s</u>ulfamate (Sal-AMS) is a selective nanomolar inhibitor of mycobactin biosynthesis that targets the enzyme MbtA,^1, 13-15^ responsible for the first and committed biosynthetic step of the mycobactins. This compound is active against whole-cell *Mtb* and in a murine model of acute TB infection but is limited by unfavorable drug disposition properties, including rapid clearance.^16-19^ To address this problem, a series of mono- and difluoro substituted furanosyl analogs were synthesized and evaluated using a combination of biological assays and NMR conformational analysis techniques.^1^ While the 2’- and 3’-monoflurination offered no improvements to both activity and pharmacokinetic properties the 2’,3’-difluorodideoxyribose derivative displayed significant gains in half-life (25-fold) and AUC (33-fold) when compared to Sal-AMS as shown in figure 1B. The improved pharmacokinetic properties, however, come at the cost in binding affinity that is shown to drop by approximately 20-fold. This is most likely due to the failure of this analog to adopt the C3’-endo (North) conformation that is the favored sugar pucker of Sal-AMS in the MbtA binding site. This hypothesis is further supported by the increase in activity noted for the 2’,3’-difluroxylose derivative. In this case the gauche and anomeric effects synergize to “ lock” the pucker angle in the North and heavily favor the C3’-endo conformation and preferred binding geometry of MbtA.

One of the primary challenges to exploring the relationship between pucker angle, biological activity, and pharmacokinetic parameters in the context of selective fluorination of the sugar moiety is the synthesis. The selective installation of fluoro groups, especially di- and tri-substitutions can be extremely laborious and in some cases, synthetically insurmountable leaving large holes in structure-activity relationship (SAR) plans and strategies. While the influence of 2’- and 3’-monofluoro substitutions on sugar conformation is well established, the effect of multiple substitutions on preference to adopt a more North or South pucker angle is more difficult to predict. It is simply impossible to gauge the balance in gauche and anomeric effects that drive the conformational equilibria when multiple electronegative groups are present. Although ab initio molecular orbital calculations have been applied in prior studies to gain insight to the intramolecular forces that determine sugar pucker angles,^10, 11^ methods of this type only provide static pictures and relative energies of model structures, typically in vacuo.^20, 21^ These methods are also not easily applied to evaluate enzyme-ligand interactions and equilibrium properties (including free energies) of ligands in solution and the bound state.^22^ Historically, these limitations have been overcome using molecular mechanics and molecular dynamics calculations that rely on empirical force fields to model the system using pairwise energy functions.^23-26^ While this technique is capable of capturing the effects of chemical substitutions on the structure and energetics of enzyme-ligand complexes the results are known to depend heavily on the parameterization of the force field.^22^ For protein or nucleic acid simulations of standard amino acids, bases, and sugars, the parameter sets are well vetted and produce excellent static and dynamic physical properties of molecules on all scales.^27-30^ The development of parameters for new entities that are not validated as part of the self-consistent force field (i.e. non-standard molecules, residues, and fragments), however, can be problematic and requires significant effort in fitting the energetics of model systems to experimentally derived data. The process has proven to be quite difficult for highly substituted carbohydrates, including ribosyl moieties and related furanoses.^21, 22^ This is in part due to the limited structural data available to derive and validate parameter sets for these systems but more so to the inherent limitations of pairwise potential energy functions in capturing the electronic effects (i.e. GE and AE) that significantly influence the preferred sugar pucker angles.

Although a number of advances have been made to improving the applicability of force field calculations to carbohydrates, prior studies have not adequately addressed the development of transferrable parameters for modeling fluoro substituted ribosyl moieties and nucleosides. In this study, we tackle this problem using the generalized AMBER force field (GAFF) as the starting point for development. The approach taken applies well established techniques and algorithms available through the AMBER suite of programs to generate and optimize atomic charge sets and torsional constants to be consistent with the GAFF and ff14sb force fields. The parameters are initially fit and validated to reproduce ab initio molecular orbital calculated energies and sugar pucker conformations of mono- and disubstituted fluorofuranose ring systems. A complete analysis of the physical properties calculated using the standard GAFF assigned parameters and the revised parameters is also presented to gain insight to the limitations of current force field methods in modeling fluoro substituted sugars. Finally, molecular dynamics calculations are applied to model sugar pucker profiles for direct comparisons with experimentally derived structural data of the six flouro substituted Sal-AMS analogs as well as other fluorinated furanose sugars reported in the literature. The results show the revised parameter set offers significant advantages in the application of GAFF to the simulation and structural analysis of fluorinated sugar moieties and nucleosides.

## Methods

### Structure generation

To properly sample the conformations of the five-membered sugar ring, 20 generic structures covering pseudorotation angles from 0 °-360° at P=18° intervals were generated using the Altona and Sudaralingam method.^31^ The 24 test structures shown in figure 2 were constructed using the Schrodinger structure builder tool and fit to the 20 generic template ring geometries to obtain Cartesian coordinates for refinement. Each structure was subsequently optimized using the *pmemd* engine within the AMBER suite of programs. The ring atoms were initially fixed to maintain the desired pucker angle during the first 1,000 steps using the steepest-descent gradient method and relaxed to obtain the final optimized set of ring conformations for charge and torsional parameter development.

**Figure 2.**
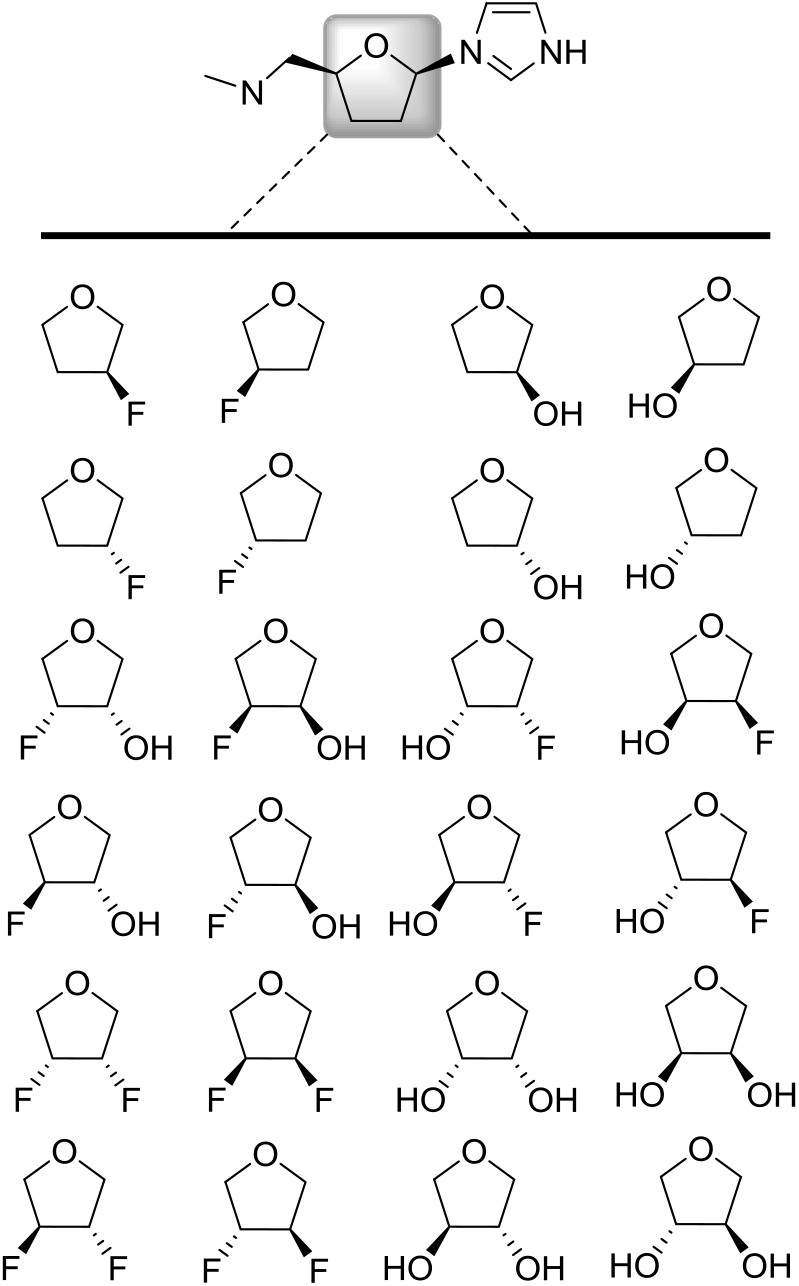
Set of 24 test (T) furanose rings used to derive partial charges and parameterize torsional varaibles.

### Calculation of new parameters

The process to calculate the new parameters was divided into two parts: i) calculation of new partial atomic charge sets for each molecule, and ii) parameterization of the torsion parameters *V*_*n*_and *γ* for the dihedrals of interest. The workflow of the process has been summarized in ***Figure 3*** and explained in detail in the following two sections along with ***Supplemental Figure 2***.

**Figure 3.**
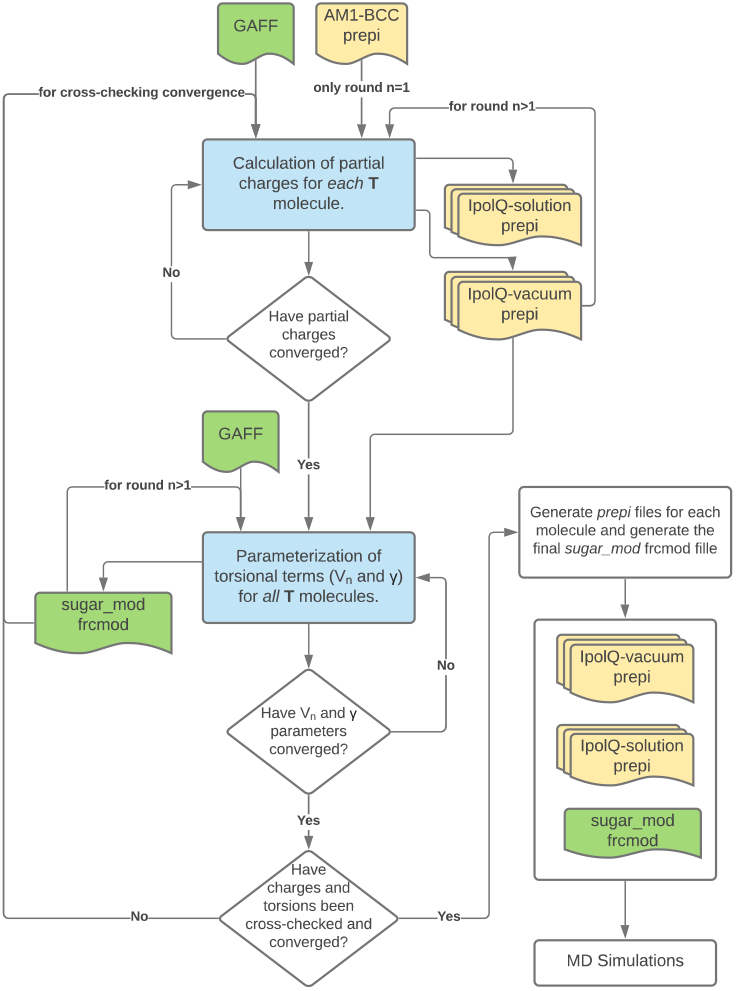
Workflow used to calculate implicitly polarized and gas-phase partial charge sets for *each* molecule, and, the *sugar_mod* frcmod file containing the reparameterization of *V*_*n*_ and *γ* for the dihedrals of interest. The processes in blue have been explained in detail in the main text sections: “ Partial atomic charges” and “ Torsional parameters” as well as ***Supplemental Figure 2***. The green and yellow files are force field and *prepi* files, respectively.

### Partial atomic charges

The IpolQ function of the mdgx program^21^ was used to generate 8 random poses for each of the 20 optimized structures spanning the pseudorotation cycle for all 24 model systems (i.e. 8 structures for each pseudorotation phase angle window of 18°). The initial set of structures was reduced using a pair-wise culling process in which any structure with a root-mean-squared-deviation less than 1Å to any other molecule was removed. The final set of structures used to calculate the partial charges was comprised of approximately 64 unique molecules for each model system. Initial charges were obtained using *antechamber* and the default AM1-BCC method. This was followed by an initial assignment of force field parameters using from the *General Amber Force Field* (*gaff*). The structures were subsequently immersed in a SPC/E water box with an 8 Å buffer zone and subjected to a minimization, heating and equilibration protocol using Amber’s *pmemd* engine (with the solute frozen in place). First, minimization was carried out using 1000 steps of steepest-descent gradient. Next, the system was heated from an initial temperature of 0K up to 300K, using an *NTV* ensemble over a total of 20,000 steps. Finally, the water molecules were equilibrated to a pressure of 1bar, using an *NPT* ensemble over a total of 30,000 steps.

Once all the subset structures had been prepared, we used Amber’s *mdgx* engine to generate submission scripts for Gaussian to calculate grids containing the electrostatic potential due to the wave function. The grids were calculated both *in vacuo* and in solution at the MP2 level using a cc-PVdZ basis set. Finally, the resulting grids were used to fit the electrostatic potential using the IPolQ procedure of *mdgx* and generate a set of partial charges for each **T**est molecule. The calculated set was compared to the previous charge set (i.e. AM1-BCC for the first iteration). If all partial charges varied less 5% from its previous value the set was determined to be converged. Otherwise, the procedure to calculate partial charges was iterated until convergence was achieved. Each new iteration used the same set of structures along with the previous IpolQ-vacuum prepi file to seed the next step of partial charges calculations. It is important to note that the AM1-BCC partial charges were only used as a starting point for the first iteration, all the following iteration steps (i.e. *i* > 2) used the previous iteration set of partial charges (i.e. *i* – 1). This procedure is summarized in ***Supplemental Figure 2***.

### Torsional parameters

The *parmchk2* utility was applied to identified all potential combinations of atoms comprising the torsional angles required to define the 24 test structures listed in ***Figure 2***. A total of 65 unique combinations were obtained (as listed in ***Supplemental Table 2***). However, since we were only interested in parameterizing those containing fluoro and hydroxy atom types only the seven shown in ***Table 2*** were fitted.

**Table 2.**
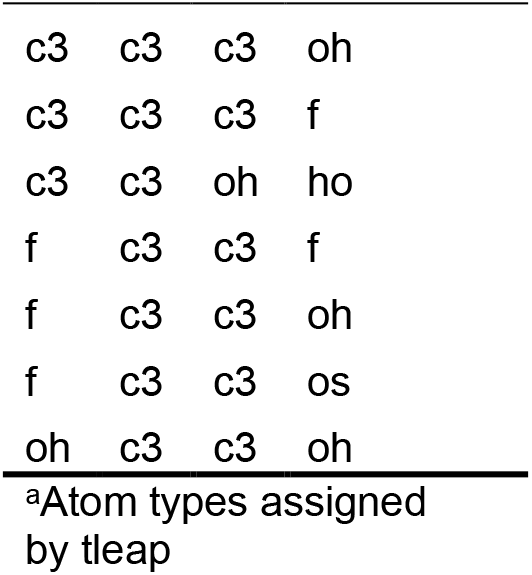
Torsion angles.^a^

|  |  |  |  |
| --- | --- | --- | --- |
| c3 | c3 | c3 | oh |
| c3 | c3 | c3 | f |
| c3 | c3 | oh | ho |
| f | c3 | c3 | f |
| f | c3 | c3 | oh |
| f | c3 | c3 | os |
| oh | c3 | c3 | oh |

<sup>a</sup>Atom types assigned by tleap

The fitting procedure was initiated by generating a total of 2480 structures for each of the 24 furanose ring templates using the *mdgx* structure generator, i.e. 124 random structures for each pseudorotation phase. Unlike the set of structures used for calculation of charge sets, none of the configurations generated at this stage were removed from the set. It has been shown that fitting a high dimensional space, i.e. multiple dihedral and angle force field parameters, requires a thorough sampling of the configurational space in order to properly parameterize the entire accessible space. For the first iteration of torsion parameter fitting, each structure (i.e. all 59,520 structures) was subsequently assigned the previously calculated *in vacuo IpolQ* charges as well as *gaff* default force field parameters. Once all the subset structures had been prepared, we used Amber’s *mdgx* engine to generate submission scripts for Gaussian to calculate single point energies for all structures at the MP2 level and cc-PVdZ basis set. Next, we used *mdgx* to calculate the residual energy, i.e. the difference between QM and MM. The residual energy, shown in ***Equation 1***, was minimized by *mdgx* through a multidimensional nonlinear least squared procedure to freely fit the value of *V*_*n*_ and a constrained value of *γ* = 0 or *γ* = 180. The remaining terms in ***Equation 1*** (force constants, bond lengths and angles, and van der Walls radii and polarizability) were not modified from the published *GAFF* values.

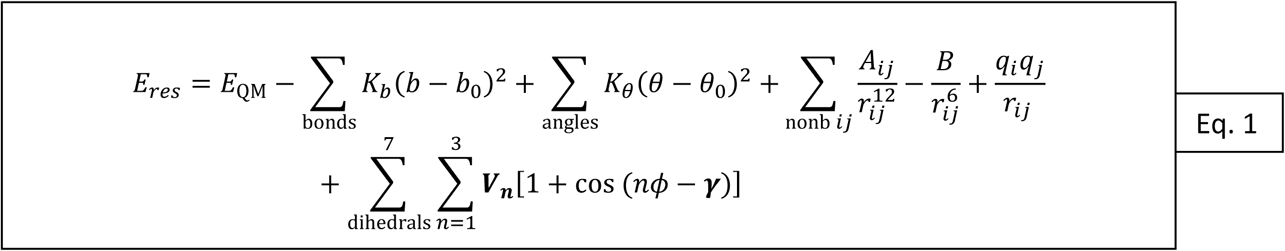

The newly fitted parameters were compared to the previous iteration force field values (i.e. *GAFF* for the first iteration). If all *V*_*n*_ values varied less 5% from its previous value **and** all *γ* remained the same, the new set of parameters were determined to be converged. Otherwise, the procedure to parameterize the fluoro and hydroxy dihedral angles was iterated until convergence was achieved. (Note: Since one of the constraints given to *mdgx* was to use a high harmonic restraint on angle stiffness constant (*arst*) the equilibrium angle (*θ*_0_) was not included in the fitting procedure.) We would like to emphasize that the original *GAFF* parameters were only used as a starting point for the first iteration, all the following iteration steps (i.e. *i* > 2) used the fitted values in the previous iteration (i.e. *i* – 1). This procedure is summarized in the flowchart shown in ***Supplemental Figure 2***. The final stage in the re-parameterization was to check that the new parameters did not change the calculated partial charge sets. Therefore, a full iteration of both partial charge set calculations and torsional parameterization was performed. Once convergence was achieved, the new parameters were assembled into a frcmod file called *sugar*_*mod*, which can be found in the supplemental information.

### Molecular dynamic simulations

All of the MD simulations were carried out using the PMEMD function of the AMBER 18 software package. All of the ligand structures were preprocessed to assign either AM1-BCC or the iteratively calculated IpolQ partial charges. The ligands were then processed using LEaP to assign *gaff* force field parameters as well as the iteratively calculated *sugar_mod* force field modification (*frcmod*) parameters. All ligands were submerged in a cubic TIP3P water box with a 10 Å buffer region. This was followed by a step-wise *NVT* heating procedure in which the system temperature was gradually ramped from 0 to 300 K over 15,000 steps and subsequently relaxed over 5,000 steps in which the average temperature was kept constant at 300 K using the weak-coupling algorithm. During both stages of heating the position of all non-solvent atoms was restrained with a harmonic potential force constant of 100 kcal/mol Å. After heating, the systems were equilibrated to a pressure of 1 bar using the *NPT* ensemble for 20,000 steps with all non-solvent atoms restrained with a harmonic potential with a force constant of 100 kcal/mol Å. This was followed by a relaxation stage. This relaxation stage varied depending whether the production run would be for: i) unrestrained simulations, or ii) simulations where the sugar ring (i.e., the five heavy atoms C1’ - C4’ and O1’) had to be constrained to a particular pseudorotation angle. In the first case, all non-solvent atoms were restrained with a harmonic potential with a force constant of 0.5 kcal/mol Å. For the second case, the heavy sugar atoms were restrained with a harmonic potential with a force constant of 100 kcal/mol Å to maintain the desired pseudorotation angle, and the rest of non-solvent atoms were restrained with a harmonic potential with a force constant of 0.5 kcal/mol Å. The minimization, heating and equilibration stages were all carried out using the PMEMD.MPI function. All production runs were carried out using the *NPT* ensemble and PMEMD.CUDA function. For unrestrained simulations the total calculation time was 200 ns per ligand. On the other hand, the phase analysis simulations required that the sugar heavy atoms be restrained with a harmonic potential with a force constant of 100 kcal/mol Å. This type of simulation was ran for 20 ns per pseudorotation angle per ligand (i.e., 360 ns per ligand).

## Results

To evaluate the ability of the sugar-mod parameter set to adequately capture the QM energy profiles of the training set test structures, a regression analysis was performed using the standard gaff force field parameter set with AM1-BCC partial charges, and the revised sugar-mod parameter set with *IpolQ* partial charges. A total of 60,000 structures were generated using the mdgx program to sample the pseudorotation space of all the molecules shown in ***Figure 2***. Structures for all molecules in ***Figure 2*** were generated to sample pucker angles at 18 degree intervals between 0 ≤ *P* ≤ 360. The MM single-point energies for each structure were calculated using pmemd with either the AM1-BCC+*GAFF* or IpolQ+*sugar_mod* parameters. The single-point QM energy was calculated, for the same set of structures as MM, using Gaussian with the B3LYP method and the CC-PVdZ basis set. The results shown in Fig.4 indicate that the *gaff* force field parameters tend to underestimate the QM calculated energy by approximately 30%, i.e. *gaff =* 0.731(QM Energy) with r^2^=0.864. On the other hand, the newly parameterized *sugar_mod* force field has an almost 1:1 relationship between MM and QM values, i.e. *sugar_mod =* 0.973 (QM Energy) with r^2^=0.976. The graph shows that the default *gaff* force field parameters underestimate the total potential energies of the ring systems, especially at higher energy values. This is an indication that energy barriers between the sugar puckering conformational phase space are not properly described by the current *gaff* parameters. As a result, transitions between minima energy wells can occur at a higher rate than their true experimental observations, leading to an erroneous characterization of the system. Our results also demonstrate that the *gaff* force field results in a broader distribution of MM energies for sugar puckering structures at QM equipotential values (see ***Supplemental Figure 1*** and ***2***, and comparison between r^2^ values). The broadening of energy sampling may become particularly problematic when calculating relative binding free energies (ΔΔG) using free energy perturbation (FEP) or thermodynamic integration (TI) methods since the error of the calculated relative free energy is directly dependent on the standard deviation of the simulation’s observed energy distribution. Consequently, a larger standard deviation results directly in larger errors and reduced confidence in calculated values.

**Figure 4.**
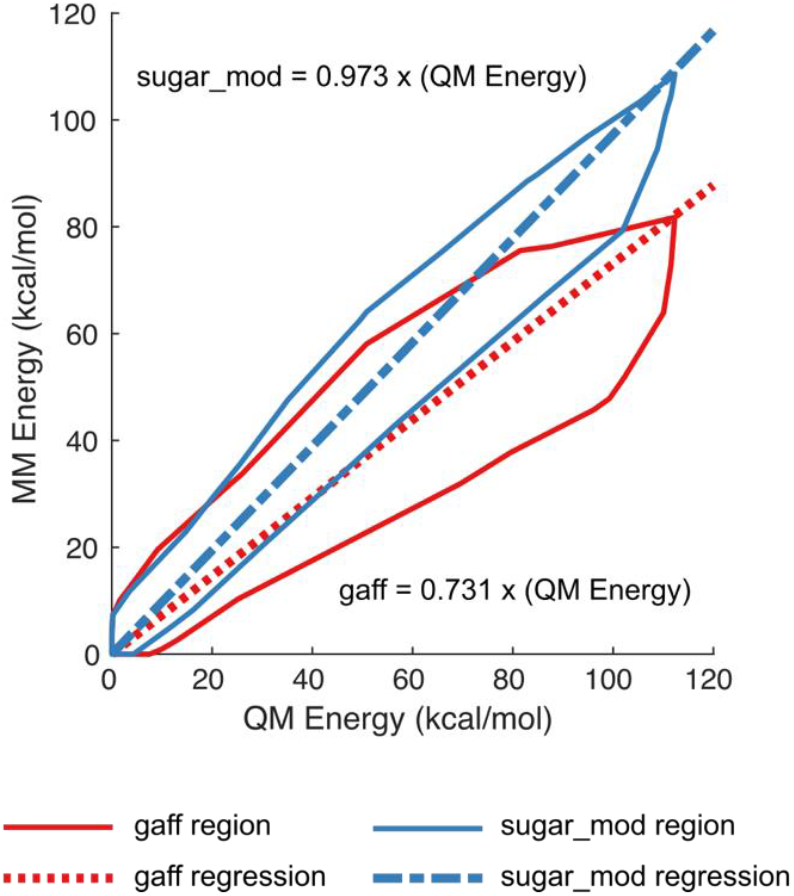
Comparison of molecular mechanics (MM) energies calculated using the *gaff* and *sugar_mod* force field parameters vs quantum mechanics (QM) energies. The total sample size used were *n* = 47596 structures. For clarity purposes only the boundary of all data points is shown rather than individual data points. The red, solid and dashed, lines represent the energies calculated using the *gaff* force field parameters. The blue, solid and dashed, lines represent the energies calculated using the *sugar_mod* force field parameters.

The *sugar_mod* force field parameter set was further evaluated using constrained MD simulations of test compounds 1-24 to sample conformational energies across the entire pseudorotation cycle. Once again, ring pucker conformations were generated at 18° intervals using the mdgx module and subsequently immersed in a periodic box of TIP3P water molecules. Positional harmonic constraints were applied to the heavy atoms of the furanose ring and all systems were equilibrated under NPT conditions (300K and 1 bar) for 20ns. The minimized QM energy for each ligand at constrained P values (EQM(P)) was calculated using the MP2 level and the cc-PvDZ basis set, and the minimized MM energy was calculated using pmemd with the IpolQ charges and *sugar_mod* parameters. The P value was constrained by freezing in place the sugar ring heavy atoms. The average energy error (EE) between the MM and QM energies for all ligands, shown in ***Figure 5***, was calculated using Equations 2 and 3, where the subscript L denotes the energy error specific to each individual test model.

**Figure 5.**
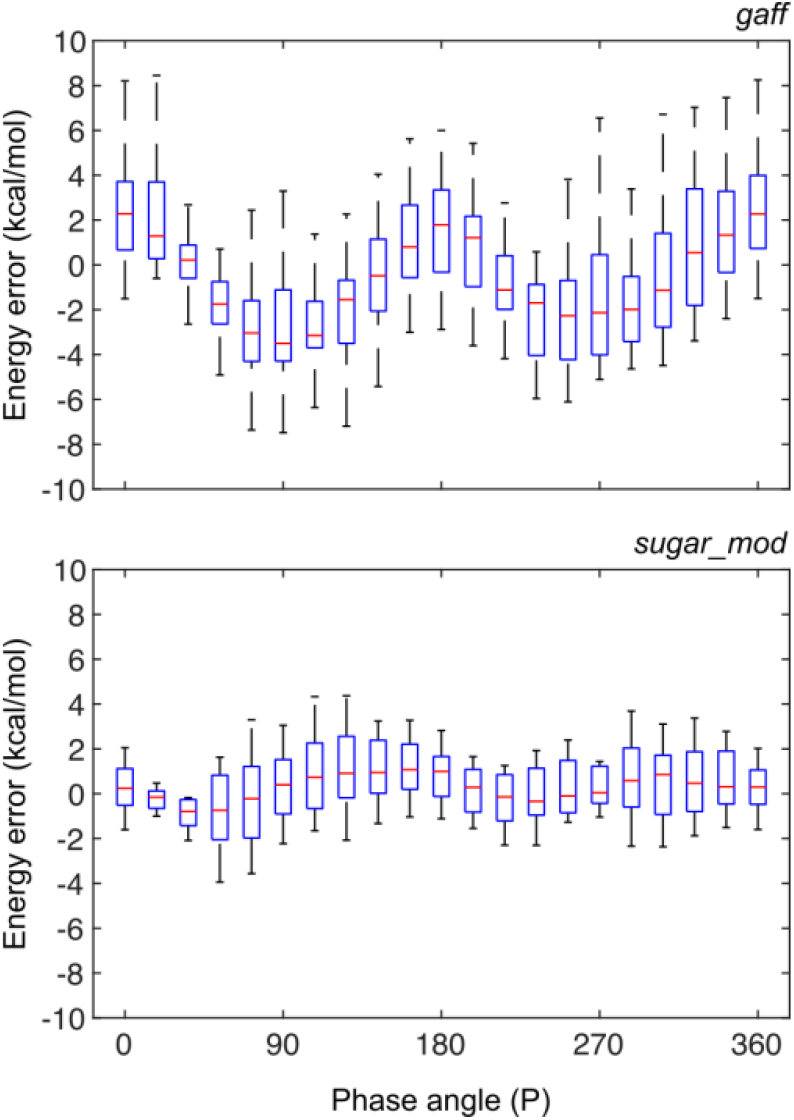
Energy error statistics vs. sugar pucker phase angle (P) for (*top*) *gaff* force field and (*bottom*) *sugar_mod* force field parameters. The energy error was calculated as the difference between the average MM energy in 20ns simulation per P value and the QM energy; both MM and QM simulations had structures with frozen sugar heavy atoms. The boxplots show the statistical analysis for structures **1**-**24**. On each distribution box, the red central mark indicates the average value (μ), the bottom and top edges of the box indicate one standard deviation (μ ± σ) and the whiskers extend to two standard deviations (μ ± 2σ).

The average energy error at a two standard deviation level (i.e. 95% of the total population) for the *gaff* force field yields an error of up to ±8 kcal/mol vs. the ±4 kcal/mol from the *sugar_mod* force field. A similar improvement is observed at the one standard deviation level, with errors of ±4 kcal/mol and ±2 kcal/mol for each respective force field. The symmetry breaks for the average error values when using the *gaff* force field (−4.3 < μ < 2.3 kcal/mol) but not for the *sugar_mod* force field (μ = ±0.9 kcal/mol). This further suggests that the *gaff* force field tends to underestimate the energy of a C2’/C3’ *mono* or *di* substituted sugar molecule. In ***Supplemental Figure 3*** we present the statistical energy average for each P value for compounds **1**-**24** with compounds parameterized using the *gaff* and *sugar_mod* force fields.

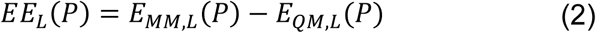

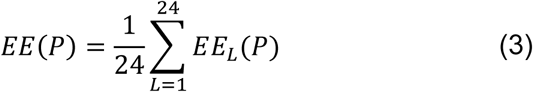

As a final measure of applicability of the new parameter set to modeling mono- and difluorinated furanose ring systems, the pucker angles and corresponding energies of compounds **2-7** were evaluated using MD simulations. Most experiments involving sugar-containing molecules report the ring configuration using the single parameter percent north (%N) or percent south (%S). However, the sugar ring system is a dynamic equilibrium between all possible pseudorotation and amplitude angles.^11^ Therefore, we simulated the behavior of the six fluoro and hydroxylated analogues of Sal-AMS (**2**-**7**) shown in ***Figure 1*** through 200 ns of unrestrained MD simulations using both the *gaff* and the *sugar_mod* force field parameters. The calculated probabilities, shown in ***Figure 6a***, were derived based on the simulation time spent at each ring puckering configuration angle. The data show that there is a remarkable difference between the configurations obtained using each of the two different force field parameters.

**Figure 6.**
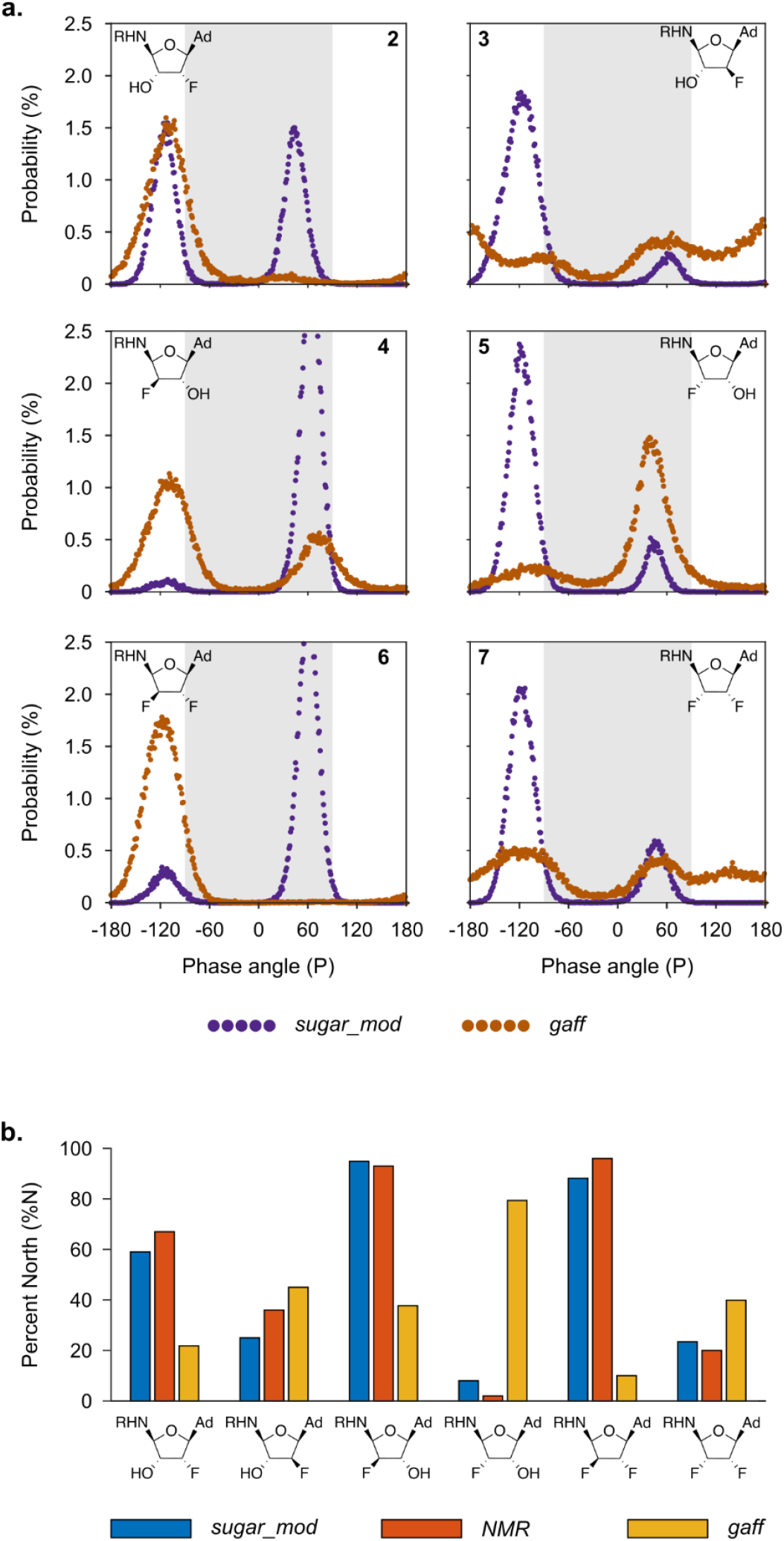
Comparison of puckering between *gaff* and *sugar_mod* force fields. **a**) Percent probability of phase angles for six fluorinated and hydroxylated Sal-AMS analogues calculated from unrestrained MD simulations using the IpolQ+*sugar_mod* (purple) and AM1-BCC+*gaff* parameters (brown). The light grey shaded region shows the “ north” configuration region, i.e. −90 ≤ P ≤ 90. **b**) Comparison of total percent north (%N) probability for six fluorinated and hydroxylated Sal-AMS analogues calculated from unrestrained MD simulations using the IpolQ+*sugar_mod* (blue) and AM1-BCC+*gaff* parameters (yellow) compared to values obtained from literature^1^ experimental NMR values.

The simulations show that the *gaff* force field parameters tend favor either one of the north (N) or south (S) configurations as shown in panels **2, 4, 5** and **6** of ***Figure 6a***. There are certain cases such as the arabino-2’,3’-OH F (**3**) and ribo-2’,3’-FF (**7**) configurations where the *gaff* results suggest there is no clear preference and there is an almost even distribution of all the puckering angles. However, NMR experimental data^1^ has demonstrated that there is a predominantly south preference for both **3** and **7**. This shows that in fact the *gaff* force field is not able to properly capture the behavior of the ring puckering for all cases and that a reparameterization of certain force field terms was warranted. On the other hand, the simulation results using the *sugar_mod* parameters show that in all the tested cases there are clear peaks at either the N or S configurations. The data obtained from the *sugar_*mod simulations show that the C2’ position of the sugar ring tends to have the greatest effect on the conformational position of the pucker. When the fluoro or hydroxyl substitution follows the 2’-endo configurations (i.e., **2, 3, 5** and **7**) the sugar pucker exhibits a southernly configuration and the 2’-exo configurations (i.e., **4** and **6**) exhibit a northerly configuration. Visual inspection of the simulations revealed that the reason for this behavior is the weak electrostatic interaction that occurs between the C2’ substituent (i.e. fluoro or hydroxyl) with an oxygen atom of the sulfamoyl moiety.

To complete the validation of the puckering parameters, we calculated the total percent north (%N) configuration, as the integration between −90 ≤ *P* ≤ 90 degrees and compared them to the calculated values from experimental NMR data.^1^ The wide range of angles chosen, as opposed to the more restricted definition that north configuration is −18 ≤ *P* ≤ 18, was due to the way that programs such as PSEUROT use NMR J-constants to calculate the %N value.^10, 11^ These programs assume there is a N/S dynamic equilibrium for the sugar ring and derives the mole fraction of the ring pucker populating *each hemisphere* of the pseudorotational cycle.^32^ The comparison between calculated %N for *gaff* and *sugar_mod* vs NMR data is shown in ***Figure 6b***.

## Conclusion

The primary goal of this work was to develop a parameter set for simulating fluoro substituted nucleosides using the tools available in the AMBER suite of programs. The approach followed standard procedures and methods to derive partial charges and torsional parameters to be consistent with the GAFF force field and standard parameter sets. Using 24 mono- and difluoro substituted furanose ring systems, the parameters were fit to reproduce ab initio molecular orbital energies of sugar pucker conformations spanning the entire pseudorotation cycle. The refinement process ensured that both high and low energy conformations were adequately sampled to reproduce the energies barriers to pucker interconversions. The results show the refined parameter set reproduced the sugar pucker energies calculated using calculated using the cc-PvDZ basis set at the MP2 level to near linearity (RMS = 0.973). The refined parameter set was also applied to predict the pucker preferences for a set of mono- and difluoro sugar substituted nucleosides for comparison with experimentally determine NMR structural data. The results show the calculated *%-Northern* pucker probabilities matched the NMR derived values within 8% error. Direct comparisons were also performed to evaluate the performance of the default gaff parameter set assigned by TLeaP. In general, the standard parameters were found to underestimate the energy barriers to pseudorotation which in turn caused the energy profiles to be flatter. This was shown to produce more variance in the pucker angles and more rapid interconversions. Essentially, the ring is too flexible. In addition, the default parameters failed to identify the dominant sugar pucker conformation of several nucleosides as shown in figure 6B. It is therefore reasonable to conclude that the default gaff assigned parameters are not suitable for modeling fluorinated sugar rings.

Given the increasing use of fluorination in the design of nucleosides and carbohydrate building blocks used in drug discovery, we expect the new parameter set should find widespread applications in the modeling both the physical properties of drug molecules. This parameter set was also developed to be consistent with the gaff force field and associated suite of programs. We purposely chose the smallest number of dihedral angles required to fit the energy profiles of the 24 test structures. The main reason for selecting only the seven parameters was to preserve as much transferability of the *gaff* force field to other substituted sugar ring systems. By keeping the same *gaff* base parameters, other types of substitutions, e.g. brominated or chlorinated compounds, would only require reparameterization of analogue dihedral angles in ***Table 1*** – for instance f-c3-c3-oh vs. br-c3-c3-oh – rather than the full set of parameters shown in ***Supplemental Table 2***. Furthermore, by selecting only 7 parameters we avoided the possibility of over-fitting the system that could have arose if all 99 angles listed in ***Supplemental Table 1*** and ***2*** were reparameterized. By overfitting the system, the sugar rings could have been “ locked” into a very small and localized pseudorotation and amplitude region, hindering the system from sampling its natural range. An over-fitted *sugar_mod* set would have shown poor configurational sampling on the opposite end of the spectrum to the under-fitted *gaff* parameters. Nonetheless, both would have same poor sampling that leads to erroneous sampling results and structural predictions.

## Acknowledgements

The authors thank the Minnesota Supercomputing Institute for a generous allocation award of computing time and software support.

## Supplemental Data

**Supplemental Table 1.**
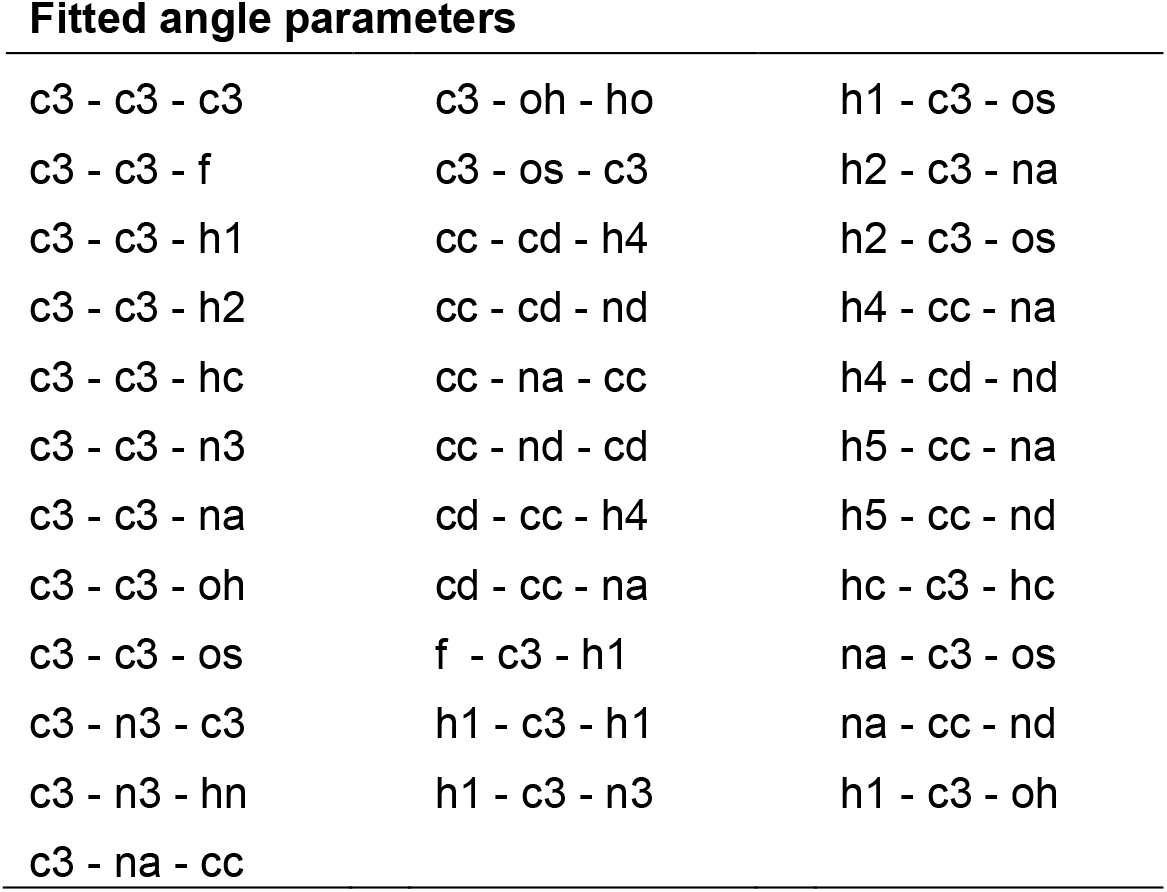
List of all angles identified by the *parmchk2* utility in the set of 24 test compounds and reparametrized using *mdgx*.

**Supplemental Table 2.**
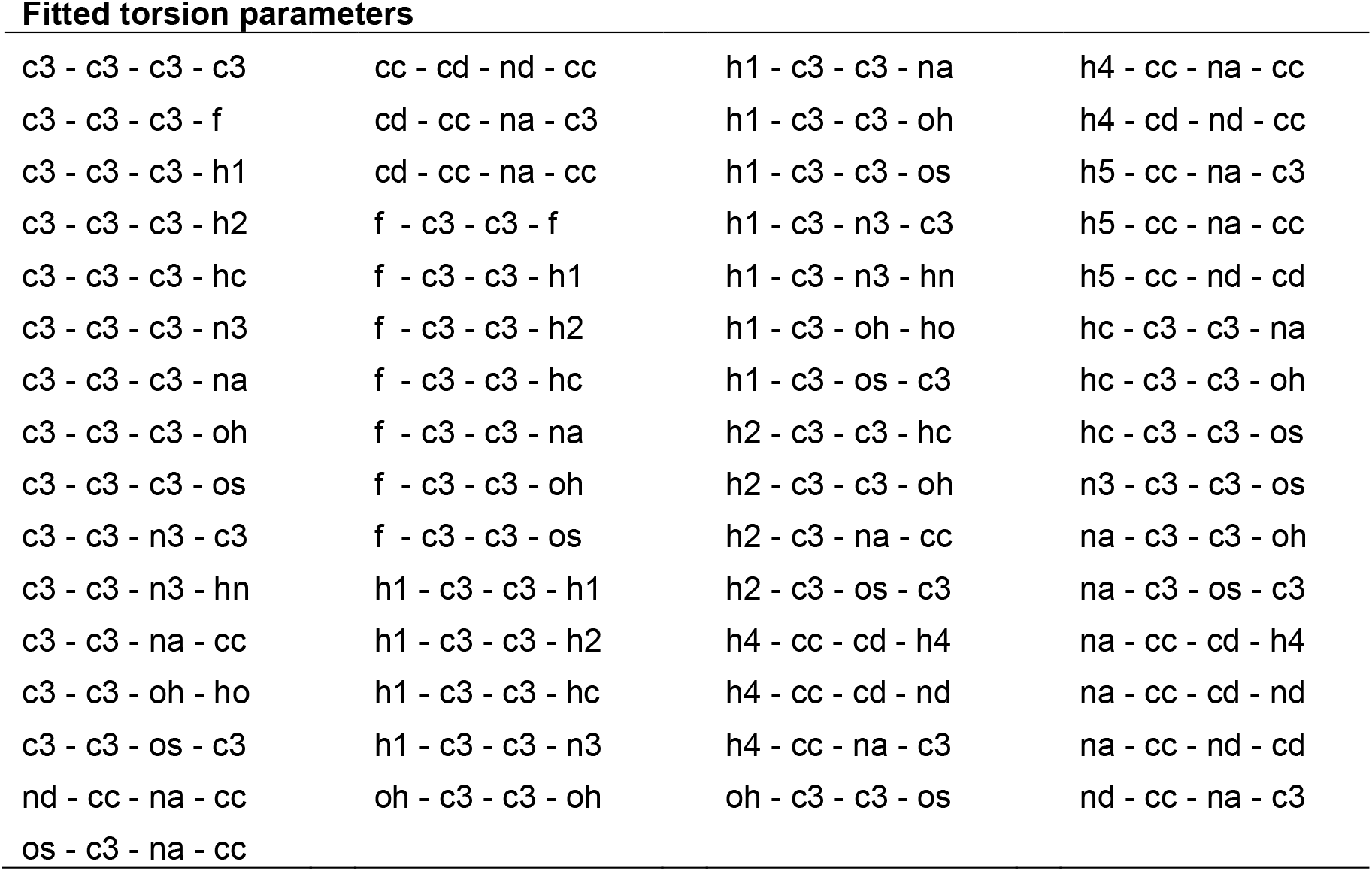
List of all torsions identified by the *parmchk2* utility in the set of 24 test compounds and reparametrized using *mdgx*.

**Supplemental Figure 1.**
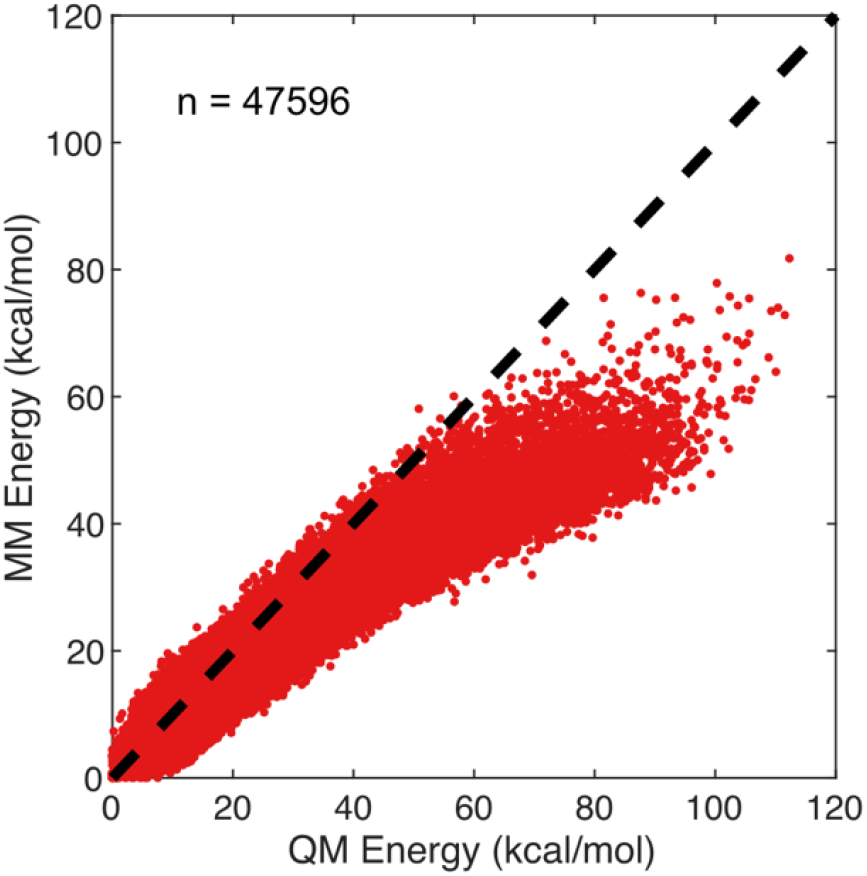
Comparison of total energy between molecular mechanics (MM) and quantum mechanics (QM) calculations. The total MM energy was calculated using the standard *gaff* force field parameters. The total QM energy was calculated at the MP2 level using the cc-PvDZ basis set. The total sample included 47596 structures, i.e. an average 1983 structures per configuration (**1**-**24**). The black dashed line indicates a 1:1 linear relationship between the two data sets.

**Supplemental Figure 2.**
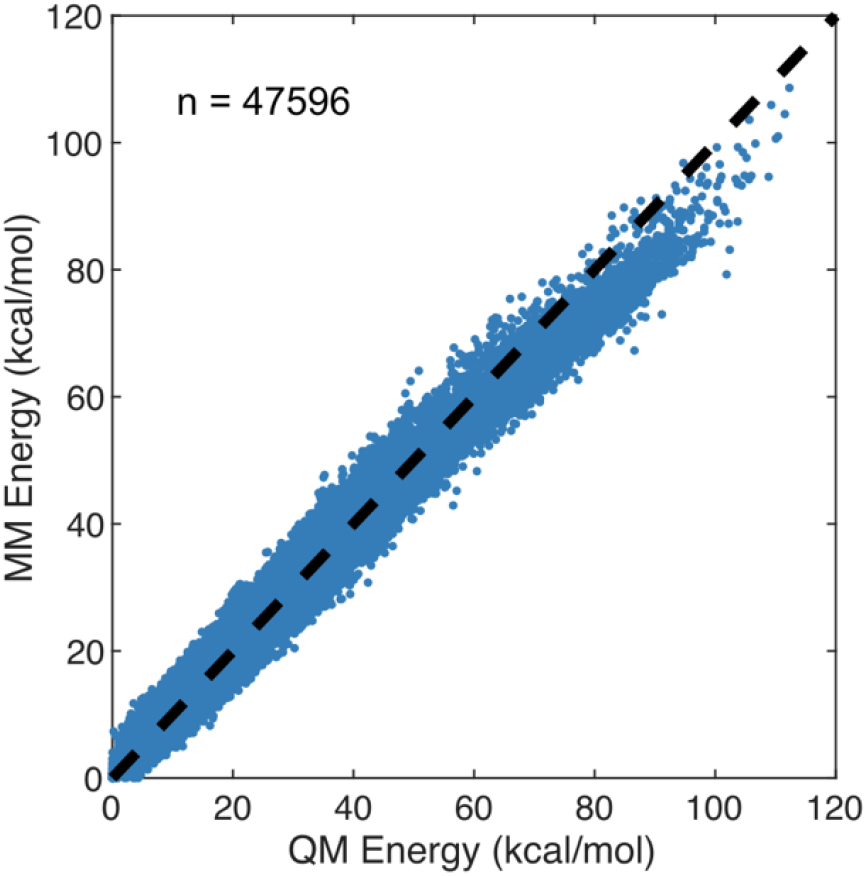
Comparison of total energy between molecular mechanics (MM) and quantum mechanics (QM) calculations. The total MM energy was calculated using the fitted parameters calculated by *mdgx*, referred to as the *sugar_mod* force field parameters. The total QM energy was calculated at the MP2 level using the cc-PvDZ basis set. The total sample included 47596 structures, i.e. an average 1983 structures per configuration (**1**-**24**). The black dashed line indicates a 1:1 linear relationship between the two data sets.

**Supplemental Figure 3.**
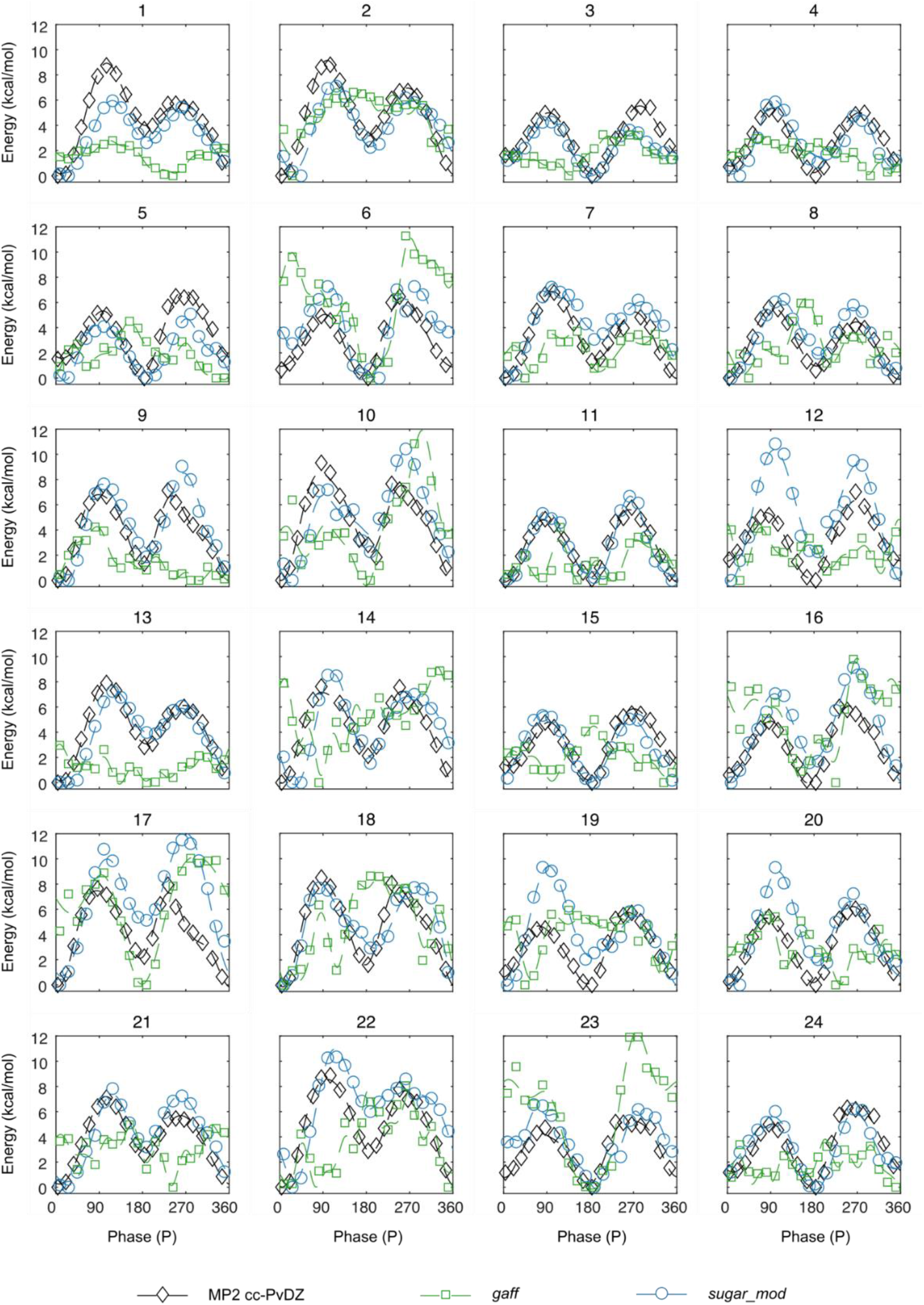
Energy profiles with sugar heavy atoms frozen at different puckering phase angles (P). Quantum mechanical energies were calculated at the MP2 level using the cc-PvDZ basis set and are shown as black diamonds. The molecular mechanical energies shown in this figure represent the statistical average for each P value during a 20ns simulation (i.e. a total simulation time of 400ns for each test structure). The results for the *gaff* and *sugar_mod* force fields are shown as green squares and blue circles, respectively.

