## Supplemental Figure and Tables for "Parameterization and Application of the General Amber Force Field to Model Fluro Substituted Furanose Moieties and Nucleosides"

### Supplemental Data

**Supplemental Table 1.** List of all angles identified by the *parmchk2* utility in the set of 24 test compounds and reparametrized using *mdgx.*

| **Fitted angle parameters** | | |
|---|---|---|
| c3 - c3 - c3 | c3 - oh - ho | h1 - c3 - os |
| c3 - c3 - f | c3 - os - c3 | h2 - c3 - na |
| c3 - c3 - h1 | cc - cd - h4 | h2 - c3 - os |
| c3 - c3 - h2 | cc - cd - nd | h4 - cc - na |
| c3 - c3 - hc | cc - na - cc | h4 - cd - nd |
| c3 - c3 - n3 | cc - nd - cd | h5 - cc - na |
| c3 - c3 - na | cd - cc - h4 | h5 - cc - nd |
| c3 - c3 - oh | cd - cc - na | hc - c3 - hc |
| c3 - c3 - os | f - c3 - h1 | na - c3 - os |
| c3 - n3 - c3 | h1 - c3 - h1 | na - cc - nd |
| c3 - n3 - hn | h1 - c3 - n3 | h1 - c3 - oh |
| c3 - na - cc | | |

**Supplemental Table 2.** List of all torsions identified by the *parmchk2* utility in the set of 24 test compounds and reparametrized using *mdgx.*

| **Fitted torsion parameters** | | | |
|---|---|---|---|
| c3 - c3 - c3 - c3 | cc - cd - nd - cc | h1 - c3 - c3 - na | h4 - cc - na - cc |
| c3 - c3 - c3 - f | cd - cc - na - c3 | h1 - c3 - c3 - oh | h4 - cd - nd - cc |
| c3 - c3 - c3 - h1 | cd - cc - na - cc | h1 - c3 - c3 - os | h5 - cc - na - c3 |
| c3 - c3 - c3 - h2 | f - c3 - c3 - f | h1 - c3 - n3 - c3 | h5 - cc - na - cc |
| c3 - c3 - c3 - hc | f - c3 - c3 - h1 | h1 - c3 - n3 - hn | h5 - cc - nd - cd |
| c3 - c3 - c3 - n3 | f - c3 - c3 - h2 | h1 - c3 - oh - ho | hc - c3 - c3 - na |
| c3 - c3 - c3 - na | f - c3 - c3 - hc | h1 - c3 - os - c3 | hc - c3 - c3 - oh |
| c3 - c3 - c3 - oh | f - c3 - c3 - na | h2 - c3 - c3 - hc | hc - c3 - c3 - os |
| c3 - c3 - c3 - os | f - c3 - c3 - oh | h2 - c3 - c3 - oh | n3 - c3 - c3 - os |
| c3 - c3 - n3 - c3 | f - c3 - c3 - os | h2 - c3 - na - cc | na - c3 - c3 - oh |
| c3 - c3 - n3 - hn | h1 - c3 - c3 - h1 | h2 - c3 - os - c3 | na - c3 - os - c3 |
| c3 - c3 - na - cc | h1 - c3 - c3 - h2 | h4 - cc - cd - h4 | na - cc - cd - h4 |
| c3 - c3 - oh - ho | h1 - c3 - c3 - hc | h4 - cc - cd - nd | na - cc - cd - nd |
| c3 - c3 - os - c3 | h1 - c3 - c3 - n3 | h4 - cc - na - c3 | na - cc - nd - cd |
| nd - cc - na - cc | oh - c3 - c3 - oh | oh - c3 - c3 - os | nd - cc - na - c3 |
| os - c3 - na - cc | | | |

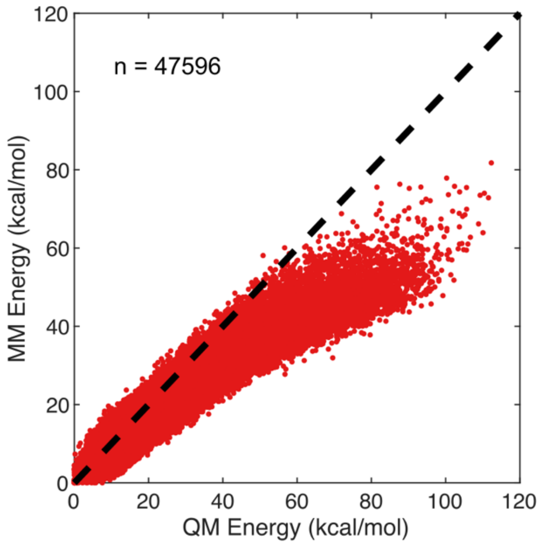


***Supplemental Figure 1.*** Comparison of total energy between molecular mechanics (MM) and quantum mechanics (QM) calculations. The total MM energy was calculated using the standard *gaff* force field parameters. The total QM energy was calculated at the MP2 level using the cc-PvDZ basis set. The total sample included 47596 structures, i.e. an average 1983 structures per configuration (**1**-**24**). The black dashed line indicates a 1:1 linear relationship between the two data sets.

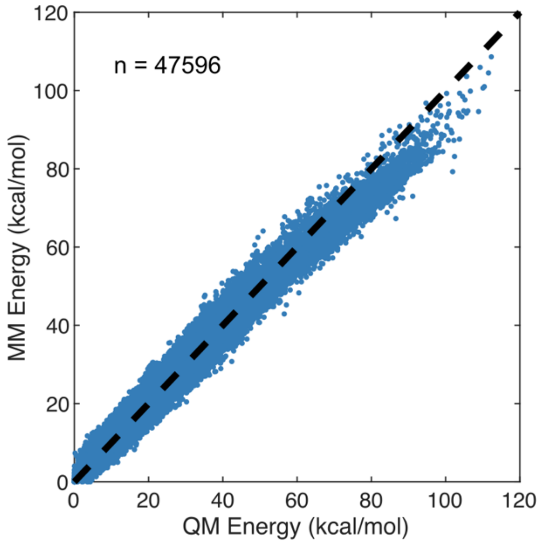


***Supplemental Figure 2.*** Comparison of total energy between molecular mechanics (MM) and quantum mechanics (QM) calculations. The total MM energy was calculated using the fitted parameters calculated by *mdgx*, referred to as the *sugar_mod* force field parameters. The total QM energy was calculated at the MP2 level using the cc-PvDZ basis set. The total sample included 47596 structures, i.e. an average 1983 structures per configuration (**1**-**24**). The black dashed line indicates a 1:1 linear relationship between the two data sets.

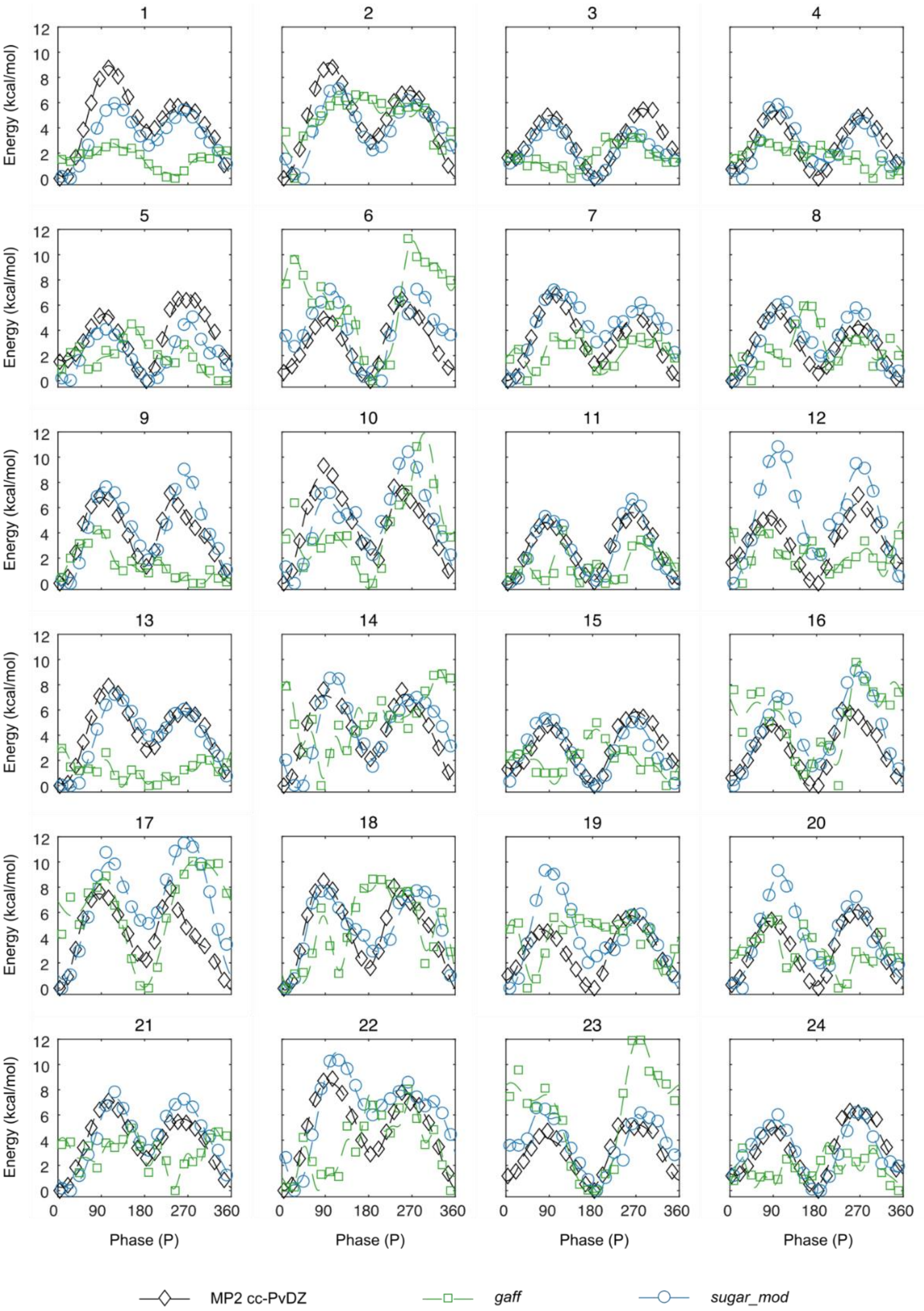


***Supplemental Figure 3.*** Energy profiles with sugar heavy atoms frozen at different puckering phase angles (P). Quantum mechanical energies were calculated at the MP2 level using the cc-PvDZ basis set and are shown as black diamonds. The molecular mechanical energies shown in this figure represent the statistical average for each P value during a 20ns simulation (i.e. a total simulation time of 400ns for each test structure). The results for the *gaff* and *sugar_mod* force fields are shown as green squares and blue circles, respectively.
